# Bicarbonate-free pH neutralization resensitizes ovarian cancer to chemotherapy and is tracked by 3T CEST MRI

**DOI:** 10.64898/2026.11.03.763942

**Authors:** Pooja A Patkulkar, Ankita Mandal, Sneha Noble, Roshni Parameswaran, Mandisa De, Shefali Jain, Shaik I Ahmed, Sanhita Sinharay

## Abstract

Ovarian tumors that initially respond to platinum-taxane chemotherapy often relapse as chemoresistant disease. The acidic tumor microenvironment is a well-known but under- exploited driver of this resistance, and neutralization strategies such as oral bicarbonate have not translated. Here we show that inhibiting specific tumor pH regulators at noncytotoxic doses relieves acidosis and restores frontline chemotherapy, monitored noninvasively at clinical 3T. In OVCAR3 and SKOV-3 ovarian cancer cells, MCT1 and MCT4 were overexpressed under acidosis, and the MCT4 inhibitor acriflavine reduced extracellular acidification at concentrations well below its cytotoxic IC_50_, without bicarbonate buffering. Acidosis blunted the efficacy of both paclitaxel and carboplatin, raising their IC_50_ values, which acriflavine reversed. In SKOV-3 tumors, acidoCEST MRI with iopromide noninvasively measured baseline tumor acidosis and its normalization following acriflavine treatment. These results establish tumor acidosis as an imageable, druggable determinant of chemoresistance and a bicarbonate-free therapeutic target.

## Introduction

A stark clinical paradox that defines high-grade ovarian cancer is an initial dramatic response to first-line platinum-taxane chemotherapy, which is then followed by a relapse with chemoresistant, ultimately fatal disease.^1,2^ Globally, 70% of ovarian cancer patients present with advanced disease (FIGO stage III–IV); where cytoreductive surgery followed by a combination of carboplatin and paclitaxel is the standard of care, which achieves high initial response rates but rarely a complete cure.^3,4^ Despite decades of added targeted and maintenance agents, this resistance persists as the central unmet need in ovarian cancer management.

A defining yet therapeutic conundrum of solid tumors is a persistently acidic extracellular microenvironment.^5,6^ Driven by glycolytic and oxidative metabolism, cells export protons and lactate through a network of pH regulators such as the monocarboxylate transporters MCT1 and MCT4, the membrane-bound carbonic anhydrase CA-IX, and the vacuolar H⁺ATPase, which maintain intracellular pH homeostasis while acidifying the surrounding interstitium.^7–10^ Far from a passive by-product of this metabolism, extracellular acidosis is increasingly recognized as an active orchestrator of the malignant phenotype and recent work identifies tumor acidosis as an environmental springboard that reprograms cancer cell energy metabolism and confers resilience to metabolic stress.^11^ Beyond promoting invasion and immune evasion, the low tumor extracellular pH (pH_e_) is also implicated in resistance to radio-, chemo and immunotherapy.^12,13^ Tumor acidosis is therefore a pleiotropic determinant of malignancy, contributing to invasion, immune escape and treatment failure alike, whose reversal has not been tested against frontline chemotherapy in ovarian cancer, or tracked noninvasively in the clinic.^14^

Weak-base chemotherapeutics such as mitoxantrone are most cytotoxic at physiological pH and lose potency as the extracellular environment acidifies.^15^ The pH dependence of paclitaxel and carboplatin is, however, suboptimally characterized, although distinct mechanisms appear to converge on a common outcome. Extracellular acidosis drives tumor cells into a reversible, more quiescent state that withdraws them from the G2/M window in which taxanes such as paclitaxel act. ^16,17^ Acidosis also impairs the cellular uptake and accumulation of platinum agents^18,19^ and diverts both drug classes into acidic lysosomal and vesicular compartments where they are sequestered and effluxed. Restoring a near-neutral tumor pH could therefore recover the intrinsic cytotoxicity of standard-of-care chemotherapy, positioning pH neutralization not solely as an adjunct, but as a determinant of drug efficacy.

Attempts to exploit this therapeutic vulnerability have primarily focused on systemic buffering, most prominently oral sodium bicarbonate, which raises tumor pH and improves outcomes in preclinical models, ^15,20,21^ but faces translational barriers, including the large, sustained doses required and poor patient tolerance.^22^ A more targeted alternative is to transiently switch off the proton and lactate export by inhibiting the proteins that act as pH regulators themselves, defined as a "pH-sensitizer" strategy that could neutralize the tumor pH at their noncytotoxic concentrations without a systemic alkaline load.^23,24^ Realizing this clinically also requires a noninvasive method of confirming that tumor pH has actually been corrected. AcidoCEST MRI reports absolute extracellular pH from the pH-dependent chemical-exchange saturation transfer of amide protons in the nonmetallic contrast agents such as Iopamidol (Isovue®) or Iopromide (Ultravist®) or Iobitridol, providing exactly such a readout. ^25,26^

Here, for the first time in ovarian cancer, we combine pH-regulator profiling, real-time metabolic phenotyping, chemosensitivity testing, and clinical-field-strength pH imaging to test whether selectively inhibiting tumor pH regulators resensitizes ovarian cancer to frontline chemotherapy. We find that MCT1 and MCT4 are overexpressed in ovarian cancer cells exposed to the acidic milieu, and that inhibition of MCT4 with acriflavine at noncytotoxic doses reverses extracellular acidification without bicarbonate buffering. We also demonstrate that acidosis confers resistance to paclitaxel and carboplatin, which acriflavine reverses; and that *in vivo*, Iopromide acidoCEST MRI at 3T noninvasively tracks tumor pH normalization and the accompanying restoration of chemotherapy response. Together, these results define an imageable, bicarbonate-free approach to resensitize ovarian cancer to standard-of-care chemotherapy.

## Results

### 1. MCT1 and MCT4 are upregulated in ovarian cancer cells under acidic conditions

To identify pH regulators that could be targeted to neutralize tumor acidosis, we profiled four principal proteins- the monocarboxylate transporters MCT1 and MCT4, carbonic anhydrase IX (CA-IX), and Vacuolar-ATPase (V-ATPase) in two ovarian cancer cell lines, OVCAR3 and SKOV-3 and in normal fallopian tube epithelial cells (hTERT-FT282). We performed immunoblotting experiments where we compared the protein expression in three different conditions – physiological pH (pH 7.4), chemically induced hypoxia, and acidic culture (pH ≤6.8) (**Fig. 1A-D**). The four regulators responded divergently to these conditions. V-ATPase, although overexpressed in the two cancer cell lines (pH 7.4) compared to normal hTERT- FT282, was not overinduced by acidosis in either line (**Fig. 1A, E**). CA-IX, a hypoxia-inducible membrane carbonic anhydrase that acidifies the extracellular space while buffering intracellular pH ^27,28^, was detected as a 54 kDa band in hypoxic OVCAR3 and as a 50 kDa species under the remaining conditions. In both cell lines studied, CA-IX was modestly higher in acidic cultures than at neutral pH, although this did not reach significance (**Fig. 1B, F**). Densitometric quantification showed that only the monocarboxylate transporters, both MCT1 and MCT4 were significantly upregulated under acidic conditions: MCT1 was ∼1.4-fold higher in acidic cultures (OVCAR3, p = 0.0009; SKOV-3, p = 0.01), and MCT4 was ∼2.5-fold higher in OVCAR3 (p = 0.001) and ∼1.3-fold in SKOV-3 (p = 0.03) (n = 3; unpaired *t*-test, 2-tailed, **Fig. 1C, D, G, H**). Although MCT1 and MCT4 both regulate lactate transport, MCT4 is the high-capacity transporter specialised for lactate efflux and is the principal protein driving extracellular acidification.^29,30^

**Figure 1.**
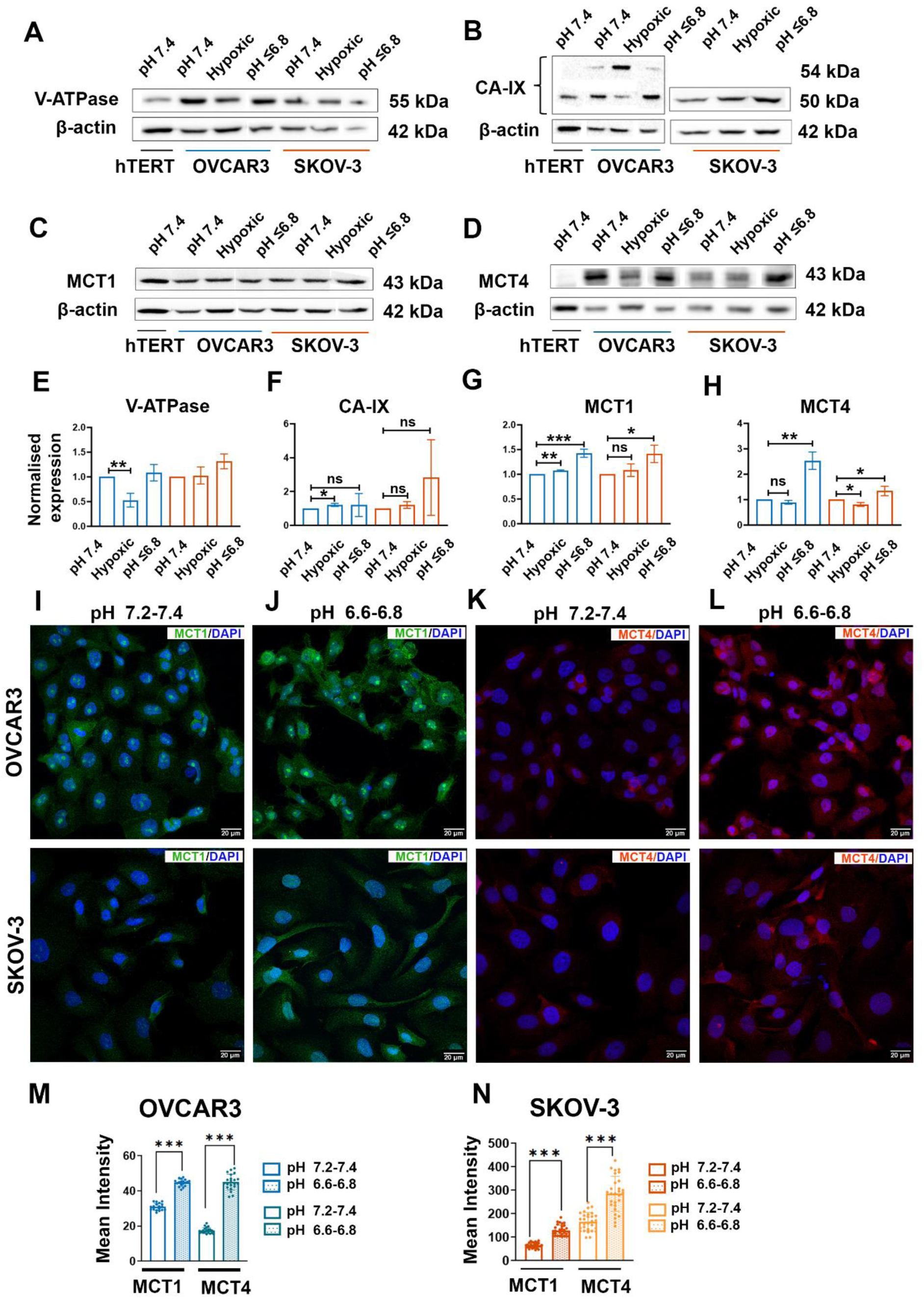
MCT1 and MCT4 are selectively upregulated in ovarian cancer cells under extracellular acidosis. (A-D) Representative immunoblots of the pH regulators (A) V-ATPase (55 kDa), (B) carbonic anhydrase IX (CA-IX; 54/50 kDa), (C) MCT1 (43 kDa) and (D) MCT4 (43 kDa) in normal fallopian tube epithelial cells (hTERT FT282, abbreviated as hTERT) and in OVCAR3 and SKOV-3 ovarian cancer cells cultured under neutral (pH 7.4), chemically induced hypoxic (CoCl₂), or acidic (pH ≤6.8) conditions. β-actin served as the loading control. (E–H) Densitometric quantification of (E) V-ATPase, (F) CA-IX, (G) MCT1 and (H) MCT4, normalized to β-actin and expressed relative to the neutral (pH 7.4) condition for OVCAR3 (blue) and SKOV-3 (orange). (I–L) Representative immunofluorescence images of MCT1 (green; I, J) and MCT4 (red; K, L) with DAPI nuclear counterstain (blue) in OVCAR3 (top row) and SKOV-3 (bottom row) cultured at neutral (pH 7.2–7.4; I, K) or acidic (pH 6.6–6.8; J, L) pH. Scale bar, 20 µm; magnification, 60X. (M, N) Quantification of mean fluorescence intensity for MCT1 and MCT4 at neutral versus acidic pH in (M) OVCAR3 and (N) SKOV-3. Data are mean ± SD, n = 3 independent experiments for E–H. Statistical significance was assessed by either unpaired t-test or one-way ANOVA with Tukey’s post-hoc analysis; ns, not significant; *p < 0.05, **p < 0.01, ***p < 0.0001.

Our immunofluorescence results in cells cultured under neutral (pH 7.2-7.4) versus acidic (pH 6.6-6.8) conditions (**Fig. 1I-L**) mirrored the same pattern observed earlier. Both MCT1 and MCT4 demonstrated significantly higher mean fluorescence intensity under acidosis in OVCAR3 and SKOV-3 (p<0.0001; **Fig. 1M, N**), corroborating the immunoblot findings. These data identified the monocarboxylate transporters, and MCT4 in particular, as the regulators most responsive to extracellular acidosis. Expression alone, however, does not establish which regulator functionally sustains acidification, and therefore we next assessed the contribution of each candidate directly, using selective inhibitors in a live metabolic flux assay.

### **2.** A small-molecule MCT4 inhibitor reduces extracellular acidification at noncytotoxic doses

Having established the transporter expression profiles under different pH conditions, we first examined the metabolic phenotype of the three lines, OVCAR3, SKOV-3 and the normal epithelial hTERT-FT282, using a Seahorse live glycolytic stress assay. Compared to normal hTERT-FT282, OVCAR3 exhibited a more pronounced glycolytic as well as energetic phenotype, whereas SKOV-3 was only slightly more glycolytic (**Fig. 2A**). The metabolic phenotype of the cell reflects the degree to which it acidifies the tumor microenvironment.^31,32^ Direct tumor-acidosis targeting approaches using sodium bicarbonate as an oral buffer established the significance of neutralization therapies in reducing aggressiveness in solid cancers.^14^ However, the translational difficulty of this simplistic approach led to exploration of other alternative strategies that exploit proton pump inhibitors (PPIs) and inhibitors of pH- transporters.^12^ Because V-ATPase inhibitors have been proposed as alkalinizing agents in solid tumors ^33,34^ we first tested the effect of the PPI, Esomeprazole (**Fig. 2B**) in lowering the extracellular pH using an induced glycolytic rate assay (Seahorse), and for both cell lines studied, we did not observe any decrease in extracellular acidification rate (ECAR) upon acute injection of the inhibitor, or the induced glycolytic proton efflux rate (induced glycoPER) at a maximum concentration of 50 µM of the PPI (**Fig. 2C, D**). This result aligned with our immunoblotting densitometry, in which V-ATPase was not significantly upregulated in either cell line under acidic conditioning, indicating that it is not an optimal target for pH-sensitization in ovarian cancer. We also examined carbonic anhydrase IX (CA-IX), with recent studies associating CA-IX expression with poor outcomes in a subset of endometrioid ovarian cancers.^35^ However, aligned with our previous observations, there was no effect of the CA-IX- specific inhibitor, SLC-0111 ^36^ (**Fig. 2B**), on ECAR and induced glycoPER for either OVCAR3 or SKOV-3 at final well concentrations of 50 µM (**Fig. 2C, D**). Next, we focused on MCT1, and MCT4, which were identified as critical players in the context of tumor acidosis and ovarian cancer in our previous experiments (**Fig. 1**). Two small-molecule inhibitors-AZD3965 for MCT1^37^ and Acriflavine hydrochloride (ACF) (**Fig. 2B)**, which impairs MCT4 function by disrupting the MCT4-basigin interaction,^38,39^ were investigated for neutralization of acidic pH_e_ through inhibition of these targets. AZD3965 produced a slight decrease in ECAR upon acute injection for both OVCAR3 and SKOV-3 at 10 µM, although the decrease was not significant (**Fig. 2C, D**). In contrast, the MCT4 inhibitor ACF, acutely suppressed extracellular acidification in both cell lines at 300 and 500 nM final concentration, and the decrease was dose-dependent (**Fig 2E, F**). A two-way ANOVA confirmed a significant effect of ACF treatment on induced glycoPER (p = 0.0009), with no significant interaction between cell line and treatment (p = 0.91). Post-hoc comparisons showed significant reductions at 500 nM ACF in both lines (OVCAR3 p = 0.010; SKOV-3 p = 0.007). This suggested that the ACF-induced increase in pH_e_ was comparable across the cell lines, even though OVCAR3 showed an absolute higher glycoPER than SKOV-3 (**Fig. 2E, F, G, H**). This comparable responsiveness is consistent with the acidosis-induced upregulation of MCT4 observed in both cell lines (**Fig. 1H, L**). In a preliminary comparison (n = 2) of ACF’s ability to lower ECAR in cells cultured in acidic versus neutral media, acidic conditions required a higher ACF concentration to increase the pH_e_, consistent with the acidosis-induced upregulation of MCT4 (**Fig 2G, H,** bottom panel). Corroborating our Seahorse ECAR and induced glycoPER results, the *in vitro* lactate assay showed that ACF treatment significantly reduced extracellular lactate efflux in both cell lines studied (two-way ANOVA, treatment effect p = 0.0003) with no significant interaction between cell line and treatment (two-way ANOVA, p = 0.25) (**Fig 2I, J**). OVCAR3 produced substantially more lactate than SKOV-3 at baseline (p < 0.0001), yet ACF significantly lowered lactate in both lines (Šídák-corrected, OVCAR3 p = 0.043; SKOV-3 p = 0.001; **Fig. 2I, J**). Together, the Seahorse and lactate assay data validate that MCT4 is a principal route of lactate export and extracellular acidification in both ovarian cancer cell lines studied, and is a critical target for pH- sensitization.^40^

**Figure 2.**
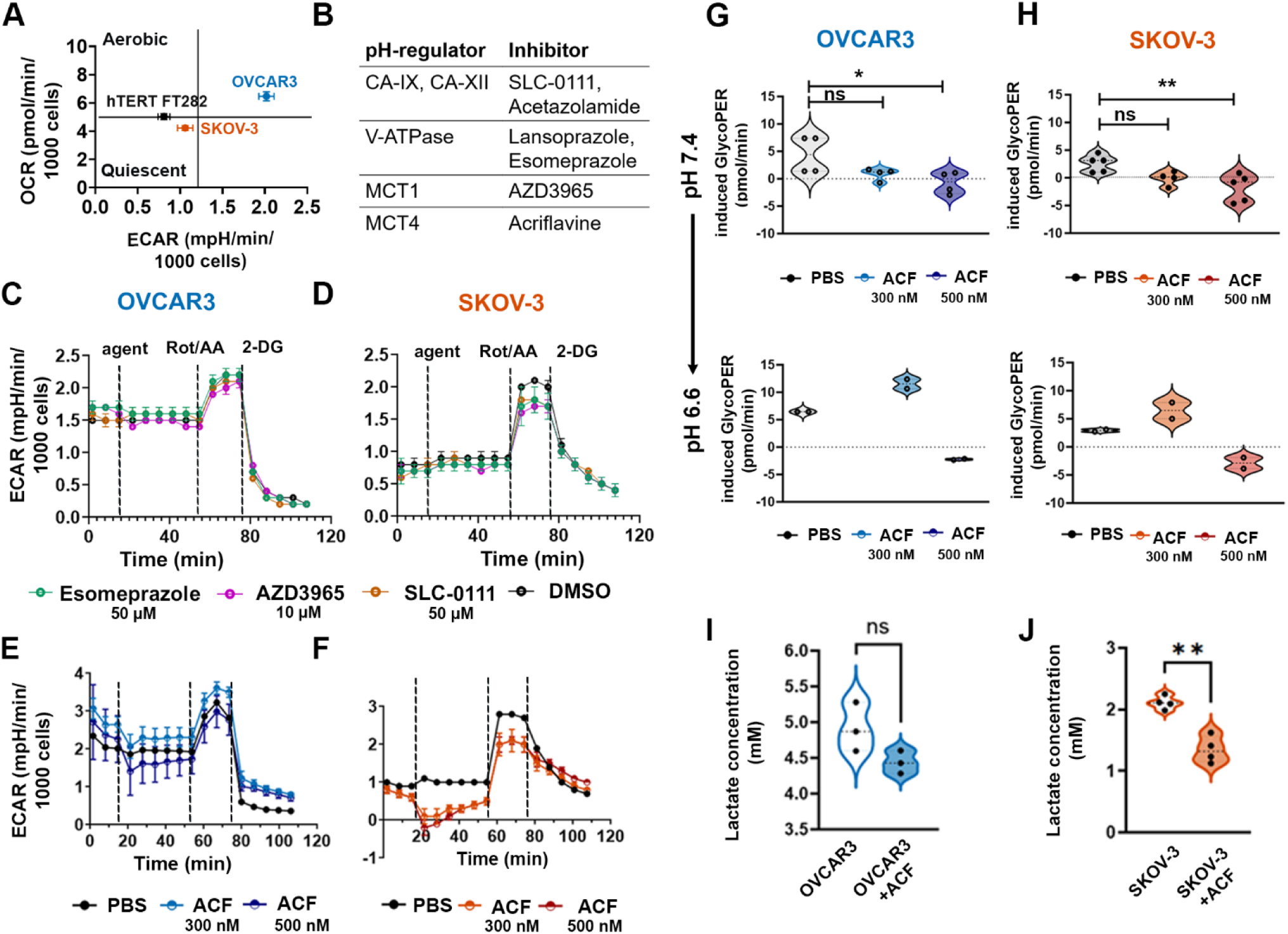
The MCT4 inhibitor acriflavine reduces extracellular acidification and lactate export. (A) Metabolic phenotype of OVCAR3, SKOV-3 and normal hTERT FT282 cells by extracellular flux analysis, plotted as oxygen consumption rate (OCR) versus extracellular acidification rate (ECAR). (B) The main pH regulators and their corresponding small-molecule inhibitors, of which four examples were tested in our studies. (C, D) Real-time ECAR traces (glycolytic rate assay) in (C) OVCAR3 and (D) SKOV-3 following acute injection of esomeprazole (V-ATPase), AZD-3965 (MCT1) or SLC-0111 (CA-IX), versus DMSO, with sequential rotenone/antimycin A (Rot/AA) and 2-deoxyglucose (2-DG) injections. (E, F) Real-time ECAR traces in (E) OVCAR3 and (F) SKOV-3 following acute injection of acriflavine (ACF; 300 or 500 nM) versus PBS, with Rot/AA and 2-DG. (G, H) Induced glycolytic proton efflux rate (induced glycoPER) in (G) OVCAR3 and (H) SKOV-3 with ACF (300 or 500 nM) versus PBS. Top, cells cultured at neutral pH 7.4 (n=3); bottom, preliminary comparison in cells cultured in acidic medium (pH 6.6; n = 2, no statistics). At pH 7.4, ACF significantly reduced glycoPER at 500 nM in both lines. (I, J) Extracellular lactate concentration in (I) OVCAR3 and (J) SKOV-3 cells treated with vehicle (control) or acriflavine (ACF). Data represented as mean ± SD; n = 3 independent experiments (A, C–F, I, J) and n = 2 (G, H bottom). Lactate and ECAR data were analysed by two-way ANOVA with cell line and treatment as factors, followed by Šídák’s multiple comparisons test. ns, not significant; *p < 0.05, **p < 0.01, ***p < 0.0001

For the small-molecule inhibitor to solely function as a pH-sensitizer rather than a co- cytotoxic agent, it should lower the extracellular acidosis without itself killing tumor cells, so that a subsequent improvement in chemotherapy response reflects pH normalization rather than added cytotoxicity. This distinction is important, as the inhibitors we examined are not merely metabolic probes, but bioactive drugs that have demonstrated their own cytotoxic effects in different doses and combinations. AZD3965 (MCT1 inhibitor, **Fig 3B**) has advanced to clinical trials as a single agent^40^, and ACF (**Fig. 3A**) has documented anti-angiogenic and antiproliferative activity.^41,42^ Each therefore carries an intrinsic cytotoxic dose range distinct from the concentration required to modulate tumor pH. We therefore first defined the cytotoxicity of ACF and AZD3965 in both OVCAR3 and SKOV-3 cell lines. The ACF IC_50_ was 4.1 µM in OVCAR3 and 1.1 µM in SKOV-3, which is respectively ∼8-fold and ∼ 2-fold above the 500 nM concentration used for pH modulation in the metabolic assays (**Fig. 3C, H**). Similarly, AZD3965 IC_50_ was 0.6 mM in OVCAR3 and 0.9 mM in SKOV-3, which is respectively ∼60-fold and ∼ 90-fold above the 10 µM concentration used for pH modulation in the metabolic assays (**Fig 3E, J**). Visually, Calcein AM staining was consistent with preserved viability of both cells at 500 nM ACF (**Fig. 3D, I**) and 10 µM AZD3965 (**Fig. 3F, K**), which was comparable to untreated cells (**Fig 3G, L),** serving as control, validating that neither the acidification screen nor the sensitizing dose acted through overt cytotoxicity. Thus, at the concentrations we employed, ACF impaired MCT4 function, and neutralized extracellular acidosis while remaining noncytotoxic, thereby separating its pH-modulating function from its known cytotoxic activity and establishing MCT4 as the effective target for further evaluation.

**Figure 3.**
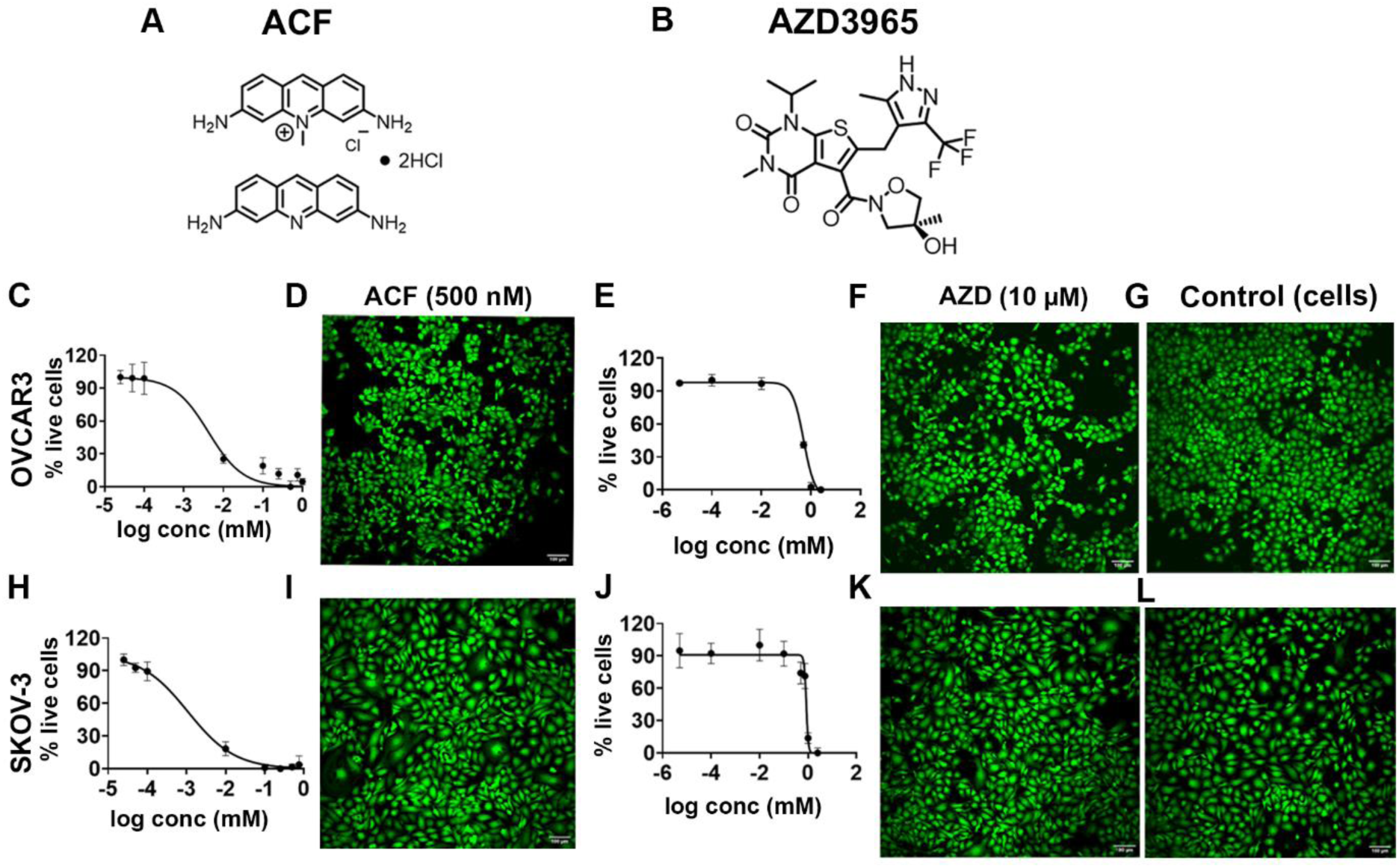
Acriflavine and AZD-3965 are noncytotoxic at the concentrations used for pH modulation. (A, B) Chemical structures of (A) acriflavine (ACF) and (B) AZD3965. (C, H) Dose-response curves (% live cells by SRB (Sulphorhodamine B assay) for acriflavine in (C) OVCAR3 and (H) SKOV-3, giving cytotoxicity IC₅₀ values of 4.1 µM and 1.1 µM, respectively. (D, I) Dose-response curves for AZD3965 in (D) OVCAR-3 and (I) SKOV-3, with IC₅₀ values of 0.6 mM and 0.9 mM, respectively. (D, F, H, I, K) Representative Calcein-AM live-cell staining (green) of OVCAR3 (top) and SKOV-3 (bottom) treated with (G, L) vehicle (control), (D) acriflavine (500 nM), or (F) AZD-3965 (10 µM). Scale bar, 100 µm. Data are mean ± SD. IC₅₀ values were calculated from nonlinear regression using GraphPad Prism 11.

### **3.** Tumor acidosis impairs paclitaxel and carboplatin efficacy, reversed by acriflavine in OVCAR3 and SKOV-3

Having demonstrated that ACF neutralizes extracellular acidosis at noncytotoxic concentrations, we investigated whether neutralizing the acidosis, restored efficacy of frontline chemotherapy used in ovarian cancer. First, we studied the dose-response of paclitaxel (1.0 nM to 750 µM) and carboplatin (0.05 µM to 5 mM) in the absence of ACF in OVCAR3 and SKOV- 3 cells cultured in the same two pH conditions (7.2-7.4 as neutral and 6.6-6.8 as acidic). IC_50_ values were obtained by fitting each biological replicate independently and reported as geometric means.

Extracellular acidosis raised the IC_50_ of the two drugs in both cell lines (**Fig. 4A-D**). The shift was most consistent for carboplatin- in SKOV-3, the IC_50_ rose from 176 µM to 706 µM (4.0-fold; 95% CI 1.6-10.4; p = 0.024; n = 3), and OVCAR3 showed a comparable 2.2- fold shift from 31.1 µM to 67.4 µM (**Fig 4E, G**). For paclitaxel, the IC_50_ also increased 19-fold in SKOV-3 (3.0 to 57.9 µM) and 3.0-fold in OVCAR3 (3.8 to 11.5 µM), but variability between independent experiments at neutral pH was substantial, and neither shift reached statistical significance (p = 0.12 and p = 0.49, respectively) (**Fig. 4F, H**). Tumor acidosis therefore reduces the potency of both components of the standard-of-care regimen, with the effect most reproducible for the platinum-based chemotherapeutic, carboplatin.

**Figure 4.**
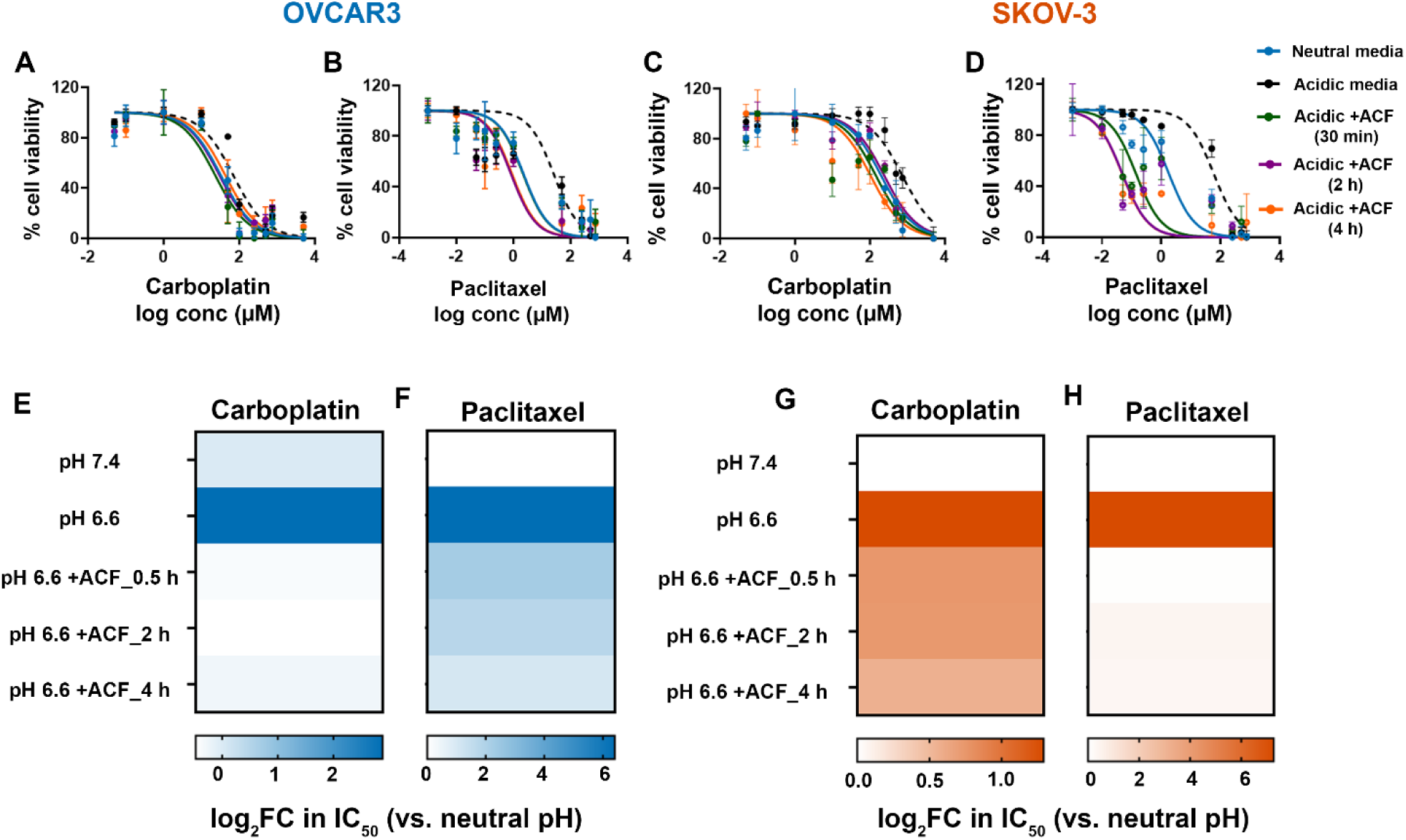
Tumor acidosis raises the IC₅₀ of carboplatin and paclitaxel, and acriflavine reverses this resistance. (A-D) Carboplatin and paclitaxel dose-response curves (% cell viability, Sulforhodamine B assay) in (A, B) OVCAR3 and (C, D) SKOV-3 cultured in neutral media (pH 7.2–7.4), acidic media (pH 6.6– 6.8), or acidic media with acriflavine (ACF) added 30 min, 2 h, or 4 h before paclitaxel. (C–F) Heatmaps of the log₂ fold-change in IC₅₀ relative to neutral pH for (E) carboplatin in OVCAR3, (F) paclitaxel in OVCAR3, (G) carboplatin in SKOV-3, and (H) paclitaxel in SKOV-3, across conditions: neutral (pH 7.4), acidic (pH 6.6), and acidic + ACF (0.5, 2, 4 h pre-incubation with paclitaxel or carboplatin).

We next asked whether ACF could reverse this resistance. ACF (500 nM) was added to cells cultured at pH 6.6-6.8, followed by paclitaxel or carboplatin 30 min, 2 h or 4 h later. ACF significantly reversed the carboplatin shift in SKOV-3 (p = 0.026–0.052 across the pre- incubation times), restoring carboplatin potency to a value statistically indistinguishable from the neutral-pH baseline (p = 0.67, 0.73 and 0.14 for 30 min, 2 h and 4 h, respectively) (**Fig 4C, G)**. The same pattern held in OVCAR3, where ACF returned the carboplatin IC_50_ to 0.71–1.21 times its neutral-pH value (**Fig. 4A, E**). For paclitaxel, ACF lowered the IC_50_ below the acidic- pH value in both lines; in some cases below the neutral-pH baseline; however, the basis for this apparent potentiation was not determined (**Fig. 4B, F, D, H**). The extent of sensitization was similar across the 30 min, 2 h, and 4 h pre-incubations of ACF, indicating the effect was independent of pre-incubation time. Together, these results show that inhibiting MCT4 with ACF reverses acidosis-induced carboplatin resistance in both cell lines, restoring drug potency to neutral-pH levels.

### **4.** Acriflavine resensitizes SKOV-3 tumors to chemotherapy *in vivo*

Building on these *in vitro* findings, we next asked whether ACF-induced neutralization of tumor acidosis restores chemotherapy response *in vivo*. Because tumor burden at the onset of treatment is itself a determinant of chemotherapy response, we first characterized the unperturbed growth kinetics of SKOV-3 xenografts in athymic nude mice, using an untreated cohort followed upto day 40 post implantation and euthanized once tumors reached 400-450 mm^3^ (n = 9, **Fig. 5A**). From this curve, we defined two pre-specified treatment initiation thresholds- T1 (when tumor size reached 350 mm^3^) and T2 (when tumor size reached 200 mm^3^) representing established and early therapeutic disease respectively (**Fig. 5A**). Overall, treatment was initiated when a tumor reached its designated threshold rather than at a fixed day post-implantation, so that every animal entered therapy at an equivalent disease burden irrespective of individual growth rate.

**Figure 5.**
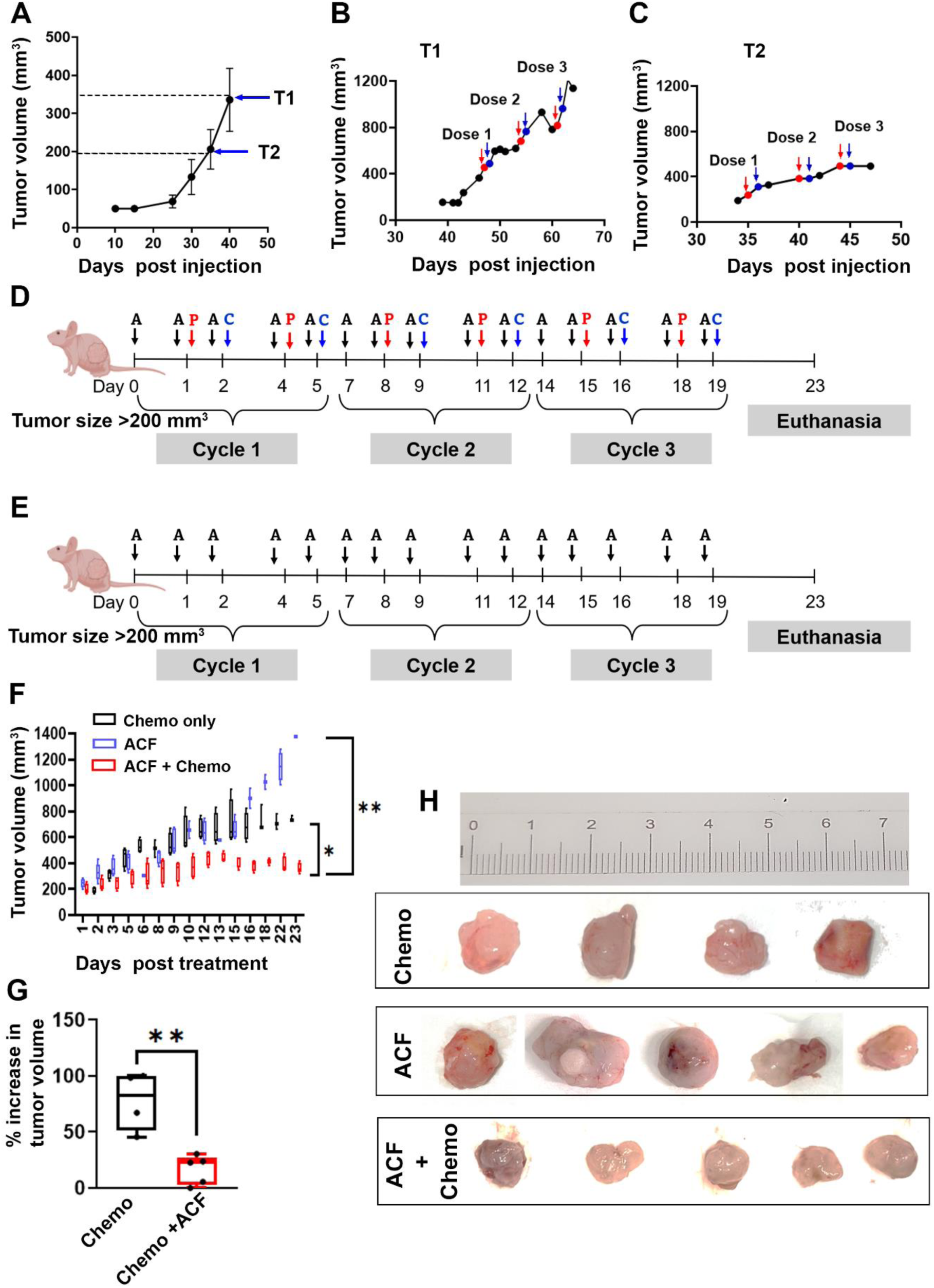
Acriflavine resensitizes SKOV-3 tumors to frontline chemotherapy in vivo. (A) Growth of subcutaneous SKOV-3 xenografts; treatment was initiated once tumors reached the therapeutic window (T1, ∼350 mm³; T2, ∼200 mm³; dashed lines). (B, C) Representative per-tumor growth in the (B) T1 and (C) T2 cohorts, showing the three chemotherapy cycles (Dose 1–3; paclitaxel, red arrows; carboplatin, blue arrows). (D, E) Treatment schedules for the (D) chemotherapy-plus- acriflavine and (E) acriflavine-alone arms, begun when tumors exceeded 200 mm³: acriflavine (A; 5 mg/kg, i.p.), paclitaxel (P) and carboplatin (C) administered over three cycles, with euthanasia on day 23. (F) Tumor growth under chemotherapy alone (P+C), acriflavine alone (ACF), or the combination (ACF+Chemo). (G) Percentage increase in tumor volume over the treatment period for Chemo (P+C) versus Chemo +ACF. (H) Representative excised tumors at endpoint: Chemo (n = 4), ACF (n = 4), and ACF + Chemo (n = 5); ruler in cm. Data are represented as box-and-whisker plot (F, G). Statistical significance by one-way ANOVA, with Tukey’s post-hoc analysis (F) and two-tailed Student’s t-test (G); *p < 0.05, **p < 0.01

Mice were assigned to three treatment groups for analysis: chemotherapy alone, (three cycles of paclitaxel and carboplatin n = 8), ACF alone (5 mg/Kg, i.p, n = 5), or ACF combined with chemotherapy (ACF+Chemo, n=5). In the combination arm, ACF dose preceded the corresponding paclitaxel and carboplatin doses within a three-cycle schedule, spanning days 0- 19, with euthanasia at day 23 (**Fig. 5D**). The ACF alone arm received an identical ACF dose and schedule without chemotherapy (**Fig. 5E**), so that this control was matched for timing as well as dose and any effect of the combination could not be attributed to ACF scheduling alone. Frontline chemotherapy alone failed to control tumor growth at either initiation threshold. Tumors treated with chemotherapy alone continued to grow through all three dosing cycles (**Fig. 5B, C**) consistent with the intrinsic chemoresistance observed clinically where a substantial fraction of ovarian cancer patients do not respond to frontline platinum-taxane therapy.^43^ Growth was nonetheless less rapid when chemotherapy was initiated at T2 than at T1 (**Fig. 5B** versus **C)**, indicating that whatever residual benefit chemotherapy retains in this resistant model is greater at lower tumor burden, and providing the rationale for initiating all subsequent combination studies at the T2 threshold.

ACF alone had no intrinsic antitumor activity, and these grew indistinguishably from chemotherapy-alone tumors (p = 0.67, one-way ANOVA with Tukey’s post hoc test), corroborating the noncytotoxic profile established *in vitro* (**Fig 5F, H**). In contrast, the combination of ACF with chemotherapy markedly suppressed tumor growth. ACF+chemo tumors grew significantly more slowly than either chemotherapy-alone (p = 0.02) or ACF alone tumors (p = 0.001) (**Fig. 5F**; one-way ANOVA, with Tukey’s post hoc analysis), and the mean increase in tumor-volume over the treatment period fell from ∼77 % with chemotherapy alone to ∼16.5 % with ACF+chemo (**Fig. 5G**; p = 0.01, Student’s *t*-test, two-tailed). Excised tumors from the ACF+chemo group were correspondingly smaller than the tumors excised from the other treatment cohorts, consistent with the volumetric data (**Fig. 5H**). Because ACF alone did not affect growth, and chemotherapy alone was largely ineffective, the response of the combination reflected resensitization of the tumors to chemotherapy rather than an additive effect of either agent.

### **5.** Acriflavine, but not chemotherapy, reduces tumor MCT1, MCT4, and LAMP1 expression without systemic toxicity

We next asked whether, in the SKOV-3 ovarian cancer model, ACF engages through MCT4 inhibition and remodels the acidic microenvironment *in vivo*, and whether this remodeling is specific to the pH-sensitizer. To achieve this, we compared SKOV-3 tumors by immunohistochemistry (IHC) across three conditions: untreated, ACF-treated, and chemotherapy-treated (paclitaxel+ carboplatin). Untreated tumors displayed strong staining for the monocarboxylate transporters MCT1 and MCT4 and for the lysosomal marker LAMP1 (**Fig. 6A**). ACF treatment showed visibly reduced staining for all three, consistent with diminished monocarboxylate-transporter–mediated proton and lactate export and a diminution of the acidic lysosomal compartment (**Fig. 6B**). In contrast, tumors treated with our combination chemotherapy regimen alone retained high MCT1, MCT4, and LAMP1 expression, comparable to untreated tumors, indicating that this microenvironmental remodeling is driven specifically by ACF and not by cytotoxic chemotherapy (**Fig. 6C**).

**Figure 6.**
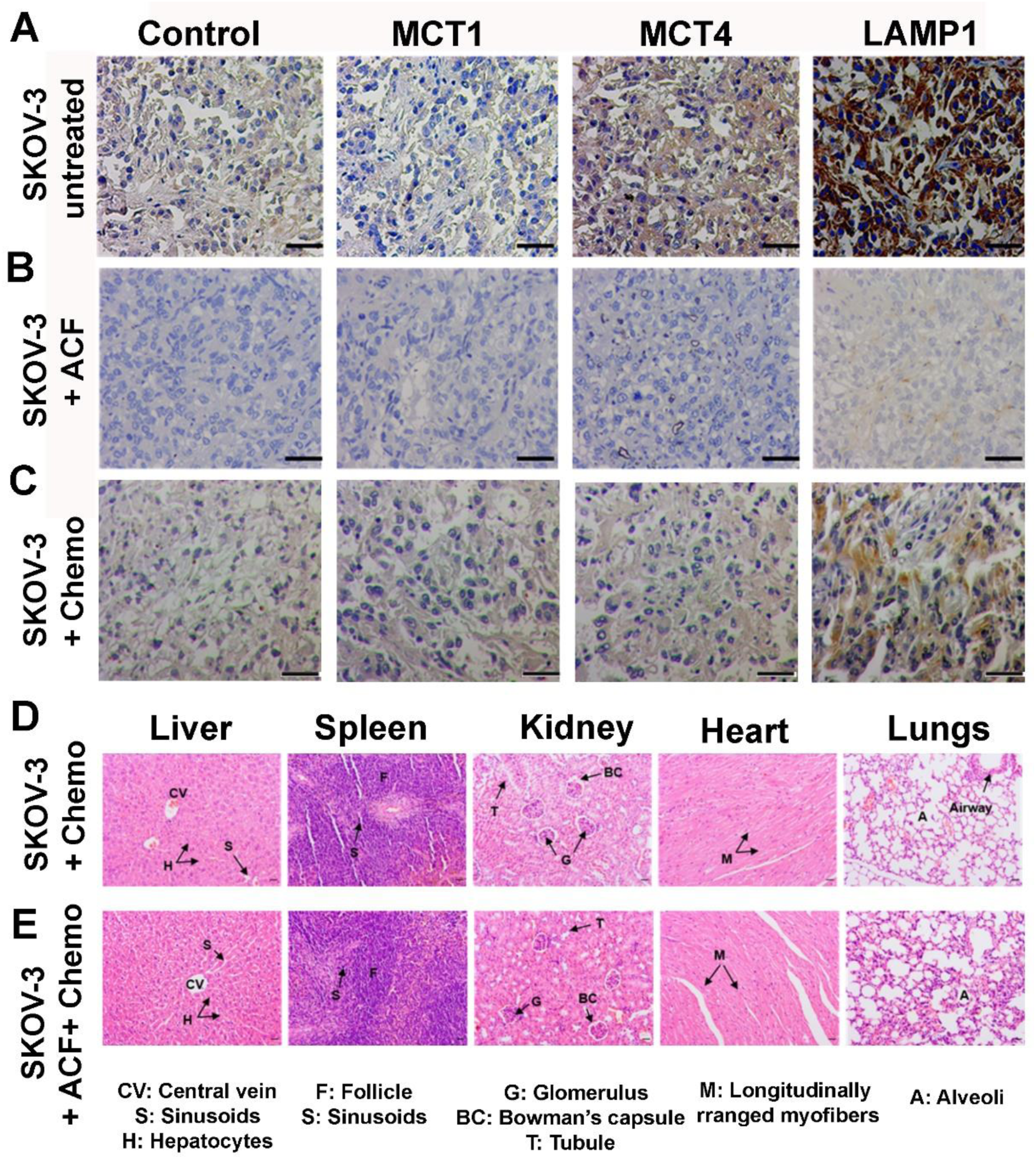
Acriflavine, but not chemotherapy, reduces tumor MCT1, MCT4 and LAMP1 expression, and the combination is well tolerated. (A-C) Immunohistochemistry for MCT1, MCT4 and the lysosomal marker LAMP1 (with a no- primary/isotype control) in SKOV-3 tumors that were (A) untreated, (B) treated with acriflavine (ACF), or (C) treated with chemotherapy (paclitaxel + carboplatin). Scale bar, 25 µm; magnification, 20×. (D, E) Representative hematoxylin & eosin (H&E) sections of major organs including liver, spleen, kidney, heart and lung from mice receiving (D) chemotherapy or (E) chemotherapy plus acriflavine. Scale bar 25 µm; magnification, 10×

Notably, LAMP1 remained high in chemotherapy-treated tumors. Because lysosomal sequestration of weakly basic and platinum drugs is an established acidosis-driven resistance mechanism, this retained acidic lysosomal compartment likely contributes to the limited efficacy of chemotherapy alone in this model (**Fig. 6C**). Mechanistically, the ACF-driven loss of LAMP1 indicates relief of this compartmental drug trapping, linking the pH normalization to restored drug action.

Finally, we assessed the tolerability of ACF combined with chemotherapy by H&E histology of major organs excised from all three cohorts. Liver, spleen, kidney, heart, and lung from mice receiving chemotherapy plus ACF retained their normal architecture, with and intact hepatic lobules (central veins, sinusoids, hepatocytes), splenic follicles, renal glomeruli, cardiac myofibers, and no substantial macrophage infiltration visible in lungs (**Fig. 6D**). the morphology was indistinguishable from that of chemotherapy-alone tumors, indicating no detectable additional organ toxicity from the addition of ACF.

### **6.** Iopromide acidoCEST MRI noninvasively maps ACF-induced tumor pH normalization in SKOV-3

*In vitro*, ACF reduced extracellular acidification and lactate efflux (**Fig. 2**), and reduced tumor LAMP1 immunostaining corroborated its effect on the acidic microenvironment (**Fig. 6A-C**), and yet the decisive test of whether ACF neutralized tumor acidosis within an *in vivo* model requires measuring extracellular pH (pH_e_) in the intact tumor, noninvasively, on a clinical platform. We therefore used CEST MRI with the clinically approved iodinated contrast agent iopromide (Ultravist®) to map tumor pH_e_ in SKOV-3 xenografts at a 3T MRI (**Fig. 7**). Iopromide carries two exchangeable amide (N-H) pools whose CEST MR signal appears at 4.4 and 5.6 ppm, referenced to water at 0 ppm. The two amide pools exchange at different rates and therefore show differential pH dependence; the ratio of their CEST effects cancels the concentration term while retaining pH sensitivity, yielding absolute pH_e_ independent of agent concentration in the tumor.^44–46^ Untreated SKOV-3 tumor-bearing mice (Cohort 1, **Fig. 7A**) that received no ACF or chemotherapeutic drugs (n = 5) were markedly acidic (**Fig. 7B, H**). Voxel-wise pH_e_ maps derived through Bloch-McConnell fitting of the two amide resonances,^26,47^ along with baseline and B_0_ correction, overlaid on anatomical T2-weighted images showed a predominantly acidic tumor interior, with mean tumor pH_e_ of 6.1 ± 0.1 (**Fig. 7G**), substantially below physiological pH and consistent with the extracellular acidosis expected of an acidic solid tumor phenotype.

**Figure 7.**
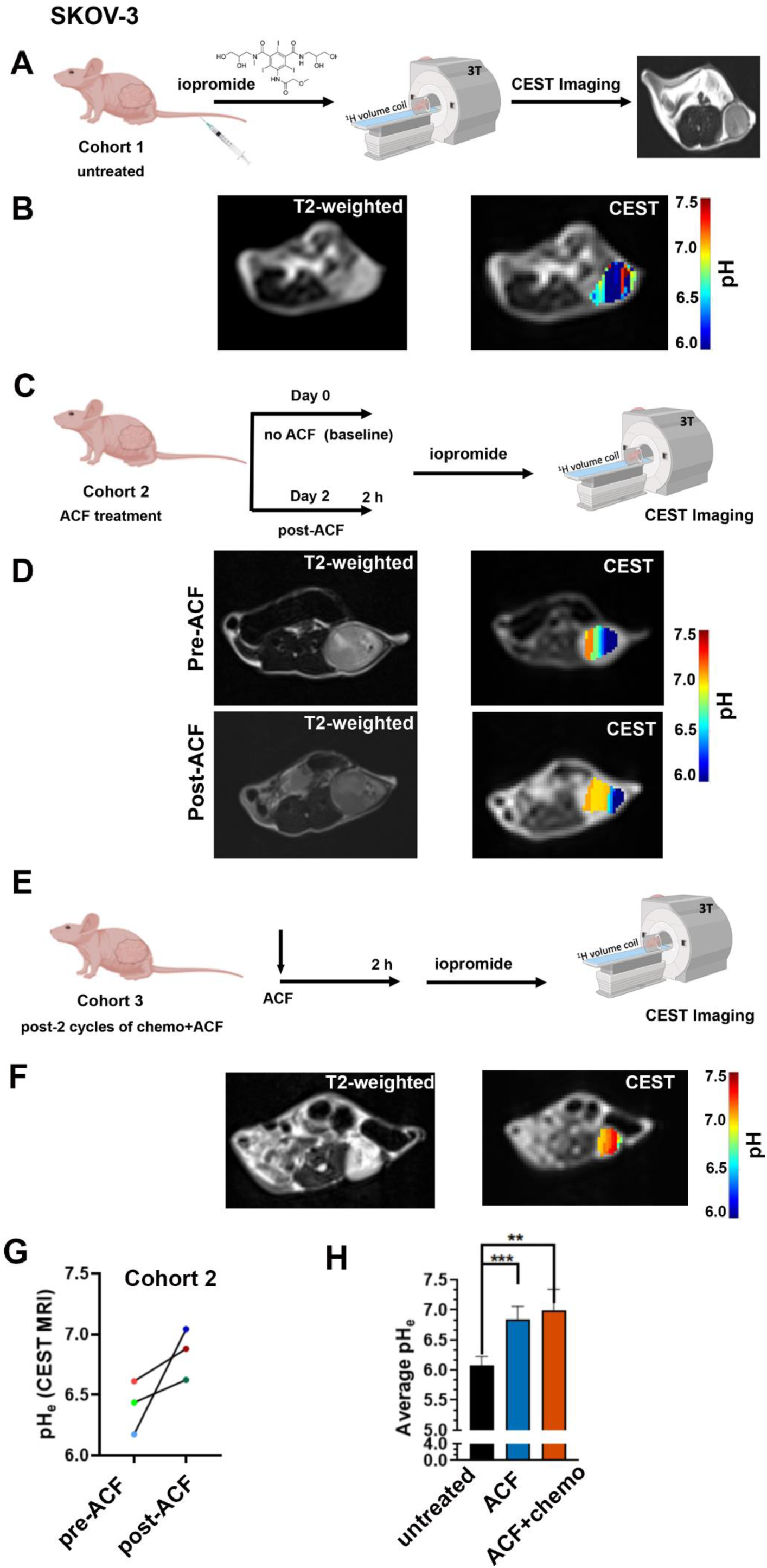
acidoCEST MRI at clinical 3T reports tumor acidosis and its neutralization by acriflavine in SKOV-3 xenografts. (A) Experimental schematic for Cohort 1 (untreated). Mice bearing subcutaneous SKOV-3 xenografts received iopromide and were imaged by CEST MRI on a 3T clinical scanner using a ¹H 32 mm volume coil (Rapid Biomedical). (B) Representative T2-weighted anatomical image (left) and corresponding acidoCEST pH map (right) of an untreated SKOV-3 tumor. (C) Schematic for Cohort 2 (ACF treatment). Each animal was imaged at baseline (Day 0, no ACF) and again on Day 2, 2 h after ACF administration, with iopromide given before each imaging session. (D) Representative T2-weighted images (left) and acidoCEST pH maps (right) from the same animal before (top, Pre-ACF) and after (bottom, Post-ACF. (E) Schematic for Cohort 3 (imaged after two cycles of chemotherapy plus ACF). Animals received acriflavine followed 2 h later by iopromide and CEST imaging at 3T. (F) Representative T2-weighted image (left) and acidoCEST pH map (right) from Cohort 3. (G). Paired tumor pH_e_ measured by acidoCEST MRI in individual Cohort 2 animals before (pre-ACF, Day 0) and after (post-ACF, Day 2) ACF treatment (H) Average extracellular tumor pH across cohorts: untreated (Cohort 1), ACF alone (Cohort 2) and acriflavine plus chemotherapy (Cohort 3). Data are mean ± SD. Significance was assessed by one-way ANOVA; **p < 0.01, ***p < 0.001; the comparison between ACF and ACF + chemotherapy was not significant. All pH maps are overlaid on the corresponding T2- weighted anatomical image and share the same colour scale

To assess the effect of ACF on extracellular tumor pH, a second cohort of SKOV-3 tumor-bearing mice (Cohort 2, n = 3) underwent longitudinal imaging (**Fig. 7C**) where each mouse was imaged with Iopromide on day 0 to establish baseline pH_e_, then re-imaged on day 2 after receiving ACF (5 mg/kg), and 2 h before intravenous iopromide and acidoCEST acquisition at 3T. ACF raised tumor pH_e_ toward physiological values. Following ACF administration, mean tumor pH_e_ increased from 6.4 ± 0.2 to 6.9 ± 0.2 (**Fig. 7D**, **G**). The magnitude of pH neutralization from an acidic baseline near pH_e_ 6.1 to a near-physiological value of 7.0 is consistent with the reduction in extracellular acidification measured *in vitro*, and provides the direct, noninvasive *in vivo* confirmation that ACF indeed neutralizes the acidic tumor microenvironment. Finally, a third cohort (Cohort 3, n = 4) that received ACF combined with chemotherapy (**Fig. 7E**), showed a comparable increase in tumor pH_e_ to 7.0 ± 0.2 by iopromide CEST, confirming that the pH correction is preserved when ACF is given with the combined chemotherapy regimen (**Fig. 7F, H**).

In summary, these findings show that acidoCEST MRI can directly visualize the action of the pH-sensitizer *in vivo*. Using iopromide, on a clinical 3T scanner, it noninvasively reported that untreated SKOV-3 tumors were significantly acidic (Cohort 1) compared to only ACF-treated (Cohort 2), (p = 0.005, one way ANOVA) or to ACF+chemo-treated (Cohort 3) (p = 0.0008, one-way ANOVA). Tumor acidity thus becomes an imageable biomarker of target engagement, and points toward image-guided pH-directed resensitization in the clinic.

## Discussion

Chemoresistance remains the principal cause of ovarian cancer mortality and is multifactorial, yet the contribution of the acidic tumor microenvironment has been difficult to monitor or exploit therapeutically.^48^ Here we demonstrate that extracellular acidosis is a reversible driver of chemoresistance in ovarian cancer, and that inhibiting the monocarboxylate transporter MCT4 with a small-molecule drug, ACF, neutralized tumor acidosis at noncytotoxic doses, restoring carboplatin efficacy and increasing paclitaxel potency. This pH sensitization could then be measured noninvasively *in vivo* through acidoCEST MRI at a clinical 3T field strength. Together, these results open an exciting avenue for connecting a druggable microenvironmental target, a bicarbonate-free pharmacologic strategy, and a clinically deployable imaging readout into a single, coherent approach to resensitize ovarian cancer to standard-of-care chemotherapy.

Historically, efforts to neutralize tumor acidity and resensitize the tumor for improved therapeutic outcomes have centred on systemic buffering, most prominently with oral sodium bicarbonate added to drinking water.^49^ Oral sodium bicarbonate raised tumor pH and inhibited metastasis in preclinical breast and prostate models, and improved chemotherapy response in breast cancer xenografts. However, all three clinical trials- in unresectable pancreatic cancer and in tumor related pain failed to demonstrate benefit of this approach, limited by poor palatability, hypokalaemia and GI intolerability.^22^ Buffer therapy also proved most effective against small tumors preclinically, a limitation of particular relevance to ovarian cancer which typically presents as bulky stage III/IV disease. More targeted approaches inhibiting individual pH regulators have been explored, though largely for direct cytotoxic effects and without a theranostic framework. Proton-pump inhibitors have been evaluated as chemosensitizers across several tumor types^50^ – in osteosarcoma, esomeprazole pretreatment enhanced cisplatin efficacy *in vivo* while having no effect on tumor growth on its own, with the accompanying phase II trial in 98 patients did not reach significance against historical control.^51^ In another relevant study, omeprazole was combined with paclitaxel in chemoresistant ovarian carcinoma models, where V-ATPase was presumed to be the operative chemosensitizing agent, although its action on pH-neutralization was not studied.^52^ Our data help explain this heterogeneity and offer a directive by first systematically profiling the pH regulators expressed under acidosis and then functionally screening their inhibitors, we identified the dominant acid-export route before intervening. Across ovarian cancer cells, only the monocarboxylate transporters, and MCT4 in particular, were consistently induced by acidosis, and functional inhibition of the V- ATPase (lansoprazole/esomeprazole) or CA-IX (SLC-0111) failed to reduce extracellular acidification, whereas MCT4 inhibition did as demonstrated through the Seahorse live metabolic flux assay and *in vitro* lactate assays. Targeting the dominant acid-export route in a given tumor type, rather than a uniformly assumed regulator, may therefore be necessary for effective, bicarbonate-free pH sensitization.

Several observations favor our argument that the resensitizing effect is mediated specifically by MCT4-dependent pH neutralization. In our IHC studies, the reduction in tumor MCT1, MCT4, and LAMP1 staining was driven by ACF and not by chemotherapy, indicating microenvironmental remodeling rather than a generic treatment effect; and ACF alone, at the dose used, neither slowed tumor growth nor was cytotoxic *in vitro*, decoupling its pH- modulating activity from its intrinsic pharmacology. The concurrent loss of LAMP1 is mechanistically informative, as lysosomal sequestration of weakly basic and platinum chemotherapeutics is an established, acidosis-driven mode of resistance,^53,54^ and the persistence of an acidic lysosomal compartment under chemotherapy alone, together with the limited efficacy of chemotherapy in this model is consistent with continued compartmental drug trapping, which ACF-mediated pH normalization appears to relieve. MCT4 was upregulated by acidosis in both cell lines, and ACF reduced extracellular acidification and lactate export in both, with no detectable difference in the magnitude of response between them. The strategy was equally effective in two cell lines with markedly different baseline glycolytic phenotypes and lactate output, which suggests that MCT4-directed pH sensitization is not contingent on a particular metabolic subtype which is a desirable property for a translational approach.

A distinguishing feature of this work is the theranostic design. Using iopromide, an approved iodinated contrast agent, acidoCEST MRI resolved the acidic baseline of ovarian tumors and the pH sensitization achieved through ACF effect produced a noninvasive readout on a standard 3T clinical scanner, with each animal serving as its own longitudinal control. Although acidoCEST MRI and its clinical translation are established^26^, to our knowledge this is the first demonstration that clinical-field-strength pH imaging can report the resensitization of tumors to frontline chemotherapy by a pH-sensitization therapy. Because the same acidity that drives chemoresistance can be imaged, corrected, and re-imaged, tumor pH_e_ offers a personalised pharmacodynamic biomarker to select patients likely to benefit capabilities that neither bicarbonate buffering nor prior pH-regulator strategies have paired with a clinical readout. These findings support the growing recognition that tumor acidosis is not a passive by-product of metabolism but an active determinant of the malignant phenotype^11^ placing therapy resistance within that framework and showing it to be pharmacologically reversible. They also align with the intrinsic pH-dependent mechanism of the frontline agents,^16,17^ reframing pH sensitization as a governing factor for drug efficacy rather than an adjuvant convenience.

There are certain limitations in our study- one being that the *in vivo* efficacy and imaging experiments used a single subcutaneous SKOV-3 model with modest cohort sizes and a confirmation in orthotopic, patient-derived, and platinum-resistant ovarian models would be a significant step to establish the generality of this proposed theranostic strategy. Secondly, ACF is a repurposed small molecule with recognized off-target activities, including HIF-1 inhibition. Several independent lines of evidence nonetheless support a MCT4-dependent, pH- mediated mechanism: MCT4 was consistently induced by acidosis in both cell lines studied. Inhibitors of V-ATPase and CA-IX failed to lower extracellular acidification while ACF did so, as shown by both Seahorse flux and lactate measurements. Additionally, the reduction in tumor MCT4 and LAMP1 staining was attributable to ACF rather than to chemotherapy; and the resensitizing dose was noncytotoxic *in vitro* and did not slow tumor growth on its own. Resolving the relative contributions of MCT4 blockade and HIF-1 inhibition will benefit from more selective MCT4 inhibitors as these become available.

In summary, we describe a bicarbonate-free, imaging-guided regimen that reverses acidosis-driven chemoresistance in ovarian cancer by inhibiting MCT4, with the therapeutic pH sensitization confirmed noninvasively on a clinical 3T platform. Our engineered approach leverages an approved contrast agent towards standard chemotherapy, requiring only a targeted, noncytotoxic pH-sensitizer and therefore offers a translationally tractable route to improving frontline therapy where the same measurement identifies acidic, chemoresistant tumors as well as predicts therapeutic response.

## Methods

### Cell Culture

The ovarian cancer cell lines OVCAR3 and SKOV-3, and the normal fallopian tube epithelial line hTERT-FT282 were a gift from Prof. Ramray Bhat (Department of Developmental Biology and Genetics, IISc). OVCAR3 was maintained in RPMI-1640 with 20% FBS, SKOV- 3 in McCoy’s 5A, and hTERT-FT282 in DMEM/F12, each supplemented with 10% FBS and 1% penicillin–streptomycin, at 37℃ in 5% CO₂. To assess protein expression under hypoxia, OVCAR3 and SKOV-3 cells were cultured in medium supplemented with the hypoxia-mimetic cobalt chloride (CoCl₂·6H₂O; 100 or 500 μM). Similarly, to induce acidosis, cells were cultured in bicarbonate-free medium (RPMI-1640 for OVCAR3, McCoy’s 5A for SKOV-3; HiMedia) supplemented with FBS; pH-neutral controls received 150 μL of 0.33 M sodium bicarbonate. Media were equilibrated at 37 °C in 5% CO₂ for 24–30 h to reach the target pH (acidic, 6.6– 6.8; neutral, 7.4), verified with a microelectrode pH meter (Horiba Scientific, Japan). Cells were then exposed to acidic or neutral medium for 24 and 48 h before lysate preparation.

### Lysate preparation

Cells seeded in normoxic, hypoxic, and acidic conditions were lysed using RIPA buffer containing a protease-phosphatase inhibitor cocktail (Sigma-Aldrich). Cells were scraped using a cell scraper, and the collected lysates were incubated on ice for 30 min. After incubation, lysates were sonicated for 15 sec and then centrifuged at 12000 rpm at 4 ℃ for 20 minutes. The supernatant was collected, and lysate aliquots were stored at -80 ℃ for protein estimation and other experiments.

### Protein estimation

Estimation of protein concentration was performed on lysates using the Bradford standard assay (BSA) in a regular 96-well plate. Briefly, following the addition of Bradford reagent to the wells, samples were incubated for 5 mins at room temperature, and then absorbance was taken at 595 nm using a plate reader (BioTek Synergy H1, Thermo Scientific, USA).

### Western blot

Expression of the pH regulators MCT1, MCT4, CA-IX, and V-ATPase was assessed by immunoblotting. Protein lysates (30 μg) were heat-denatured (95 °C, 5 min), resolved on a 10% gel (5% stacking), and transferred to nitrocellulose membrane using a Bio-Rad Trans- Blot Turbo system. Membranes were blocked in 5% BSA in TBST (1 h, room temperature) and incubated with primary antibody (1:1000 in 5% BSA) overnight at 4 °C with shaking. After three TBST washes, membranes were incubated with HRP-linked anti-rabbit IgG secondary antibody (CST 7074S; 1:2000 in TBST) for 1 h at room temperature, washed three times in TBST, developed with luminol–peroxide (1:1; Bio-Rad), and imaged on a Bio-Rad ChemiDoc XRS+. Bands were analyzed using the Image Lab software. For loading controls, membranes were stripped (mild stripping buffer, 45 min), re-blocked in 5% BSA (1 h), and reassessed for β-actin (1:1000, 1 h, room temperature) following the same secondary-antibody and detection steps. Primary antibodies: CA-IX (CST 5649S), MCT1/SLC16A1 (CST 85680S), V-ATPase (CST 14617S), HIF-1α (CST 36169S), and β-actin (CST 4970T), all rabbit monoclonal from Cell Signaling Technology (CST); and MCT4 (Santa Cruz sc-376140).

### Immunofluorescence assay

OVCAR3 and SKOV-3 cells were plated on glass coverslips and cultured in neutral (pH 7.0- 7.4) or acidic medium (pH 6.6-6.8) for 48 h. Cells were washed three times in PBS, fixed in 4% PFA, permeabilized with 0.5% Triton X-100, and blocked in 2% BSA (2 h, room temperature). Cells were incubated overnight at 4 °C with primary antibody diluted in 1% BSA - rabbit monoclonal anti-MCT1 (CST 76508S, 1:75) or mouse monoclonal anti-MCT4 (Santa Cruz sc-376140, 1:50) followed by the corresponding Alexa Fluor–conjugated secondary antibody (1:300, 2 h, room temperature): anti-rabbit IgG Alexa Fluor 488 (CST 4412S) for MCT1 and anti-mouse IgG Alexa Fluor 594 (CST 8889S) for MCT4. Nuclei were counterstained with DAPI. Images were acquired on an Olympus FV3000 upright confocal microscope at 60×. Mean fluorescence intensity was quantified in ImageJ from ROIs drawn around individual cells (∼20–30 cells per group) and plotted as a bar graph with individual values.

### Sulforhodamine B (SRB) Assay to determine cytotoxic range for small molecule inhibitors of the pH-regulators

The colorimetric SRB assay was performed to measure cytotoxicity of the various small- molecule inhibitors of the pH-regulators of interest. IC₅₀ values were determined for this panel of pH-regulator inhibitors spanning the major regulator classes -the MCT inhibitors acriflavine hydrochloride (MCT4) and AZD3965 (MCT1), the carbonic anhydrase inhibitor acetazolamide (pan-CA inhibitor) and SLC0111 (CA-IX inhibitor), the V-ATPase/proton-pump inhibitor esomeprazole to define a noncytotoxic sensitizing range. Cells were seeded in 96-well plates and, 24 h later, treated across a concentration range of 10 nM-5 mM for 48 h at 37 °C in 5% CO₂. Three biological replicates were performed per condition. Cells were fixed with 30% trichloroacetic acid (4 °C, 1 h), washed, and air-dried overnight. Fixed cells were stained with SRB (15 min, room temperature), washed with 1% acetic acid to remove unbound dye, and the bound dye solubilized in 10 mM Tris. Absorbance was read at 510 nm on a plate reader (BioTek Synergy H1, Thermo Scientific, USA).

### Calcein AM staining

Cells were seeded at a density of 10,000 per well in a 96-well plate and maintained in complete culture media. The following day, the cells were washed with PBS and replaced with phenol- free media without serum. Cells were then stained with 5 µM Calcein AM (Thermo Fisher, Calcein AM C1430) and incubated for 30 min at 37°C in a CO₂ incubator. Following incubation, the staining solution was replaced with phenol-free medium. Fluorescence images were acquired using a Leica SP8 confocal microscope with a GFP filter.

### Lactate Assay

The cells were maintained in phenol red-free medium with 5% FBS, then treated with 500 nM acriflavine for 24 h. After incubation for 24 h, the cellular media were collected in a tube and used as a sample for the determination of extracellular lactate concentration. For the *ex vivo* lactate assay, a tumor sample of 250 mg was suspended in 200 µL 1X PBS and sonicated for 20 min to homogenize the tumor. The homogenate was then centrifuged at 150g for 5 minutes at room temperature. After centrifugation, the homogenate is diluted 10 times with 1X PBS and kept in a dry bath at 80℃ for 30 min to heat-denature protein, especially lactate dehydrogenase (LDH). After denaturation, we centrifuged again at 150g for 5 min at room temperature and collected the clear supernatant, which we used as a sample for the lactate assay.

For the lactate assay, we followed the instructions provided in the lactate assay kit (Sigma, MAK329). We first prepared lactate solutions for the standard plot with concentrations ranging from 2 mM to 0.2 mM. For the assay, 20 µL of sample/standard solution is used, and 76 µL of a freshly prepared reaction mixture is added according to the mentioned protocol. Immediately after the addition, the initial absorbance was recorded at 565 nm. The plate was incubated for 20 min at room temperature, and the absorbance was measured post-20 min at 565 nm. Final absorbance values were subtracted from the initial absorbance to determine the sample lactate concentration using the equation obtained from the standard curve.

### Cytotoxicity assay (SRB) with chemotherapeutic drugs under normal and acidic culture conditions

OVCAR3 and SKOV-3 cells were assayed under neutral (pH 7.0-7.4) and acidic (pH 6.6-6.8) conditions. For the neutral arm and the acidic arm without any pH-sensitization, cells were pre- incubated in acidic medium for 48 h before treatment with paclitaxel at a concentration range of 750 µM-1 nM or carboplatin at a concentration range of 5 mM-50 nM across a concentration series (nM–mM); and then following incubation with either drug for 48 h at 37 °C, standard SRB assay was performed as detailed earlier.

For the acidic arm with pH-sensitization, cells in acidic medium (pH 6.6-6.8), were pre-treated with acriflavine (500 nM) for 30 min, 2 h, or 4 h before addition of paclitaxel at a concentration range of 750 µM- 1 nM or carboplatin at a concentration range of 5 mM- 50 nM, incubated for 48 h at 37 °C and standard SRB assay was performed. IC₅₀ values were determined from three biological replicates per condition.

### Seahorse induced glycolytic rate assays (GRA) for live monitoring of extracellular acidification rate changes upon pH-sensitization

Extracellular acidification rates (ECAR) were measured in live cells on an Agilent Seahorse XFp analyzer (model number S7802A). OVCAR3 and SKOV-3 cells were seeded (20,000 cells/well) in XFp miniplates in their respective complete media (RPMI-1640 + 20% FBS for OVCAR3; McCoy’s 5A + 10% FBS for SKOV-3) and incubated overnight; sensor cartridges were hydrated the preceding day per the manufacturer’s instructions. On the day of assay, medium was replaced with assay medium (pH 7.4±0.1) and cells were equilibrated in a non- CO₂ incubator (37 °C, 45 min).

To assess pH modulation, the induced glycolytic rate assay was run with the pH sensitizer (acriflavine or AZD3965) injected acutely such that the final port A concentration is 300 and 500 nM for acriflavine and 1 and 50 µM of AZD3965, followed by addition of rotenone/antimycin A (Rot/AA, port B) and 2-deoxyglucose (2-DG, port C), yielding extracellular acidification rate (ECAR) and glycolytic proton efflux rate (induced glycoPER). Compound concentrations of Rot/AA, and 2-DG followed the respective Agilent XFp user guides. The assay was performed in three biological replicates, and the data were analyzed in Wave software (Seahorse, Agilent, USA).

## Animal experiments

All animal experiments were conducted following the guidelines mandated by the Institute Animal Ethics Committee (IAEC) of the Indian Institute of Science (Protocol number CAF/ethics/039/2024). All experiments were performed on 6–8-week-old female athymic nude mice. All the mice used in this study were housed in the Central Animal Facility (CAF) at IISc till they were ready for experiments conducted on the 3T MRI. The SKOV-3 flank tumor model was developed for this study by injecting 5 × 10^6^ cells in media along with Matrigel (1:1 mixture) on the left flank of the mice. Tumor volume was measured with a vernier calliper and calculated according to the formula Volume = ½ * (width^2^ *length). SKOV-3 tumor bearing mice (n=9) that belonged to the untreated cohort, did not receive either the pH-sensitizer, ACF or any chemotherapy drug. The imaging studies and histology studies were performed for this cohort when the tumor size attained 400-450 mm^3^.

## Experimental design for therapeutic dose and timing optimization

A total of 17 animals were used to optimize the treatment plan for our study, these animals were not used in the imaging, and at end point, they were utilised for the histology and Immunohistochemistry (IHC) experiments. Paclitaxel and carboplatin, two commonly used first-line therapy drugs to treat ovarian cancer patients, are given either in combination with the pH-sensitizer acriflavine hydrochloride (ACF) or without ACF. We divided our study into different cohorts to estimate the optimized dose and timing. In cohort 1 (n=5), chemotherapy treatment with paclitaxel (20 mg/kg) and carboplatin (50 mg/kg) were administered intraperitoneally on consecutive days when the tumor size reaches 350-400 mm³. This regimen was continued every week for 3 cycles, and the animal was euthanized once tumor size exceeded 600 mm^3^. Organs and tumors are fixed in 4% PFA and sectioned (5 µm slice thickness) for IHC with markers corresponding to MCT1, MCT4 and LAMP-1. In cohort 2 (n = 3), the same combination chemotherapy treatment was administered when the tumor size reached 150-200 mm³. This is continued every 3 days for 3 cycles, and the animal was euthanized once tumor size exceeded 600 mm^3^ or 15 days post the last cycle whichever was sooner, and organs and tumor were processed in the manner described earlier. In cohort 3 (n = 5), we treated mice with only ACF (5 mg/Kg) once the tumor size reached ∼ 200 mm³ and followed this ACF administration on 5 days/week upto 25 days. The tumor size was monitored to observe the sole effect of ACF on these tumors and the excised tumors were processed for IHC markers- MCT1, MCT4 and LAMP1. In the final cohort 4 (n = 4) we initiated the pH- sensitization of the tumors with ACF (5 mg/kg) when the tumor size reached ∼200 mm³ and followed ACF administration on consecutive days till the 3-cycle combination chemotherapy dose was completed. The continued administration of ACF was performed to ensure that the tumor remained pH-sensitized for the entire chemo-treatment cycle. Organs and tumors from this cohort were also processed in a similar manner and sectioned for assessment of the same markers as mentioned earlier.

## Histology and Immunohistochemistry studies (IHC)

The SKOV-3 tumor was excised, fixed with 4% paraformaldehyde (PFA), and stored in 0.1% sodium azide solution. Samples were then given to the RVD lab for sectioning and H&E staining. Formalin-fixed, paraffin-embedded tissue sections (5 µm) were deparaffinized in xylene, rehydrated through graded ethanol, and rinsed in PBS. Endogenous peroxidase activity was quenched with 3% H_2_O_2_ for 30 mins. Antigen retrieval was performed in citrate buffer (10 mM, pH 6) at 60 ℃ for 30 min, followed by cooling to room temperature. Sections were then blocked with 2% BSA for 2 h. Slides were then incubated overnight at 4 ℃ with primary antibodies (MCT1, MCT4, LAMP1; diluted 1:100 in 1% blocking buffer), washed with PBS, and incubated with HRP-conjugated secondary antibodies (1:300 dilution) for 1h at room temperature. Visualization was achieved using 3,3′-diaminobenzidine (DAB) substrate, followed by haematoxylin counterstaining. Finally, sections were dehydrated with sequential ethanol, followed by xylene, and mounted with anti-fade mounting solution (ThermoFisher). Stained slides were imaged using an Olympus coloured microscope (model number IX73) with 20x magnification (MCT1, MCT4, LAMP1) and 10x magnification (H&E).

### AcidoCEST MRI *in vivo* studies

We performed CEST MRI studies with iopromide (acidoCEST) to determine the extracellular tumor pH (pH_e_) before and after treatment with the pH sensitizer acriflavine hydrochloride. All MRI studies were conducted in a 3T Siemens Prisma scanner equipped with a 32 mm 1H volume transmit-receive coil (Rapid Biomedical™, Germany). For the untreated SKOV-3 tumor bearing mice that received neither the pH sensitizer ACF nor chemotherapy drug, were imaged with only Iopromide to evaluate the baseline pH of these SKOV-3 tumors. Briefly, the mouse was anesthetized with 60 mg/kg Ketamine and 16 mg/kg Xylazine injected intraperitoneally using a 31G needle and then 200 μL of iopromide contrast agent was administered i. p, following which the mouse was secured to a cradle and positioned inside the scanner. Throughout the scans, the mouse was cushioned using a heat pad to maintain physiological body temperatures. A multislice T2 scan (RARE) was performed and then a 2D single slice T2-weighted image was acquired with the best tumor position, and this was selected for CEST. For all CEST scans, a train of 40 rectangular pulses with a saturation power of 1 μT, a pulse duration of 50 ms, a TR of 15 ms, a TE of 2.83 ms and an offset frequency ranging from -10 ppm to 10 ppm were used. The field of view chosen was 164x164 mm^2^ and the matrix size was 160x192 mm^2^ with a slice thickness of 4 mm, acquired at 4 averages resulting in a scan duration of 13 minutes.

In case of the cohort where we assessed the effect of ACF on tumor pH, 3 animals were longitudinally imaged, where baseline pH_e_ of the tumors were measured using CEST MRI in the manner described above. On Day 2 following that, the same mice received 5 mg/kg ACF injection 2 hours prior to the iopromide injection and then CEST data was acquired using the same parameters. For the sensitization chemotherapy cohort, we followed the treatment regimen as outlined in Fig 5D. After completion of two cycles on chemotherapy, during the third cycle where treated mice received chemotherapy drugs (20 mg/kg paclitaxel and 50 mg/kg carboplatin) along with ACF (5 mg/kg), we performed CEST scans with iopromide to evaluate the effect of sensitization chemotherapy on tumor pH_e_. The ACF and iopromide injection were at an interval of 2 h from each other on the day of CEST scan, whereas the paclitaxel and carboplatin injections on the animal were performed a day or two prior to the ACF, iopromide injections. The CEST data were analyzed using Bloch-McConnell equations with fitting parameters for pH determined using customized code in MATLAB R2024a (MathWorks).

## Statistical analysis

All statistical analyses were performed in GraphPad Prism. Unless stated otherwise, data are presented as mean ± SD, and n refers to the number of independent biological replicates; technical replicates within an experiment were averaged prior to analysis. All tests were two- tailed, and p < 0.05 was considered significant. Significance is indicated as *p < 0.05, **p < 0.01, ***p < 0.001, ****p < 0.0001; ns, not significant. For immunoblot, the densitometric values were normalized to β-actin and expressed relative to the neutral-pH (pH 7.4) condition. Comparisons were made by unpaired two-tailed *t*-test or by one-way ANOVA with Tukey’s multiple comparisons test, as indicated in the figure legends. Induced glycoPER values measured from Seahorse extracellular flux assay and extracellular lactate were analysed by two-way ANOVA with cell line and treatment as factors, followed by Šídák’s multiple comparisons test. The interaction term was used to test whether the magnitude of the response to ACF differed between OVCAR3 and SKOV-3. Dose-response curves were fitted in Prism using the log(inhibitor) versus normalized response model. Each biological replicate was fitted independently, and 95% confidence intervals were determined by profile likelihood. Because IC_50_ values are log-normally distributed, all averaging and statistical comparisons were performed on logIC_50_; and reported as geometric means. Tumor volumes were compared across treatment groups by one-way ANOVA with Tukey’s multiple comparisons test.

Tumor pH_e_ measured before and after ACF administration through acidoCEST MRI in the same animal was compared by paired two-tailed *t*-test. Average tumor pH_e_ across the untreated, ACF and ACF+chemotherapy cohorts was compared by one-way ANOVA with Tukey’s multiple comparisons test.

## Acknowledgements

SS acknowledges the Central Animal Facility (CAF), IISc, for help with intravenous injections and support for animal experiments performed in this study. SS acknowledges the JN Tata MRI Centre, IISc for support with MRI studies.

## Author contributions

SS and PP conceived the project and PP designed and performed the experiments and anlaysed the data. SS and AM and PP wrote the manuscript, and AM analysed data and performed the experiments. SN analysed the data. RP contributed to experimental design and conducting experiments. MD and AM and SJ contributed to data analysis and data interpretation. SA contributed to data analysis and code fitting.

